# Meningeal macrophages mask incision pain sensitization in male rats

**DOI:** 10.1101/2025.07.25.666793

**Authors:** Mahshad Kolahdouzan, Shahrzad Ghazisaeidi, YuShan Tu, Milind M. Muley, Eder Gambeta, Michael W. Salter

## Abstract

**Introduction:** Meninges surrounding the brain and spinal cord house a variety of immune cell types including macrophages that express the CD206 mannose receptor. Here, we investigated whether CD206^+^ macrophages in the meninges play a role in regulating nociception and pain hypersensitivity.

**Methods:** We selectively depleted CD206^+^ macrophages in the meninges around the lumbar spinal cord by intrathecal administration of anti-CD206 coupled to saporin, and determined the effects of CD206^+^ macrophage depletion on responses in naïve rats and in those that had received a skin incision to the upper hindlimb. In addition, we used RNAseq to investigate transcriptional changes in lumbar meninges and dorsal root ganglia. Experiments were done in both male and female rats.

**Results:** Depleting CD206^+^ meningeal macrophages did not alter basal responses in naïve animals of either sex. By contrast depleting these cells after skin injury induced mechanical hypersensitivity in male rats, without changes in thermal sensitivity but had no effect in females. In male rats with skin incision injury, we found that the mechanical hypersensitivity induced by depleting CD206^+^ meningeal macrophages was reversed by administering the NMDAR antagonist, APV. In addition, the hypersensitivity was reversed by an enhancer of KCC2 function, CLP290. Unexpectedly, skin incision caused significant transcriptional changes in the meninges, but only in male rats.

**Conclusions:** Taken together, our results indicate that while CD206^+^ meningeal macrophages do not regulate basal nociception in naïve rats, after skin incision injury, these cells mask mechanical hypersensitivity in male rats only. Thus, we conclude that in a sex-dependent manner CD206^+^ meningeal macrophages prevent the spread of pain hypersensitivity after a minor injury. Importantly, the skin incision we used was comparable to that used in ‘sham’ controls in numerous rodent studies of neuropathic pain. Our findings have, therefore, potentially broad implications for re-interpreting results from previous neuropathic pain research.

## Introduction

Chronic pain is a pervasive health, social, and economic problem affecting approximately 27% of the global population ^1^. Current therapeutic options remain limited due to insufficient efficacy and unacceptable side effects. Physiological pain is critical for responding to tissue-damaging or potentially tissue-damaging stimuli. However, pathological pain, such as neuropathic pain stemming from injury, lesions or disease of the nervous system or inflammatory pain stemming from tissue inflammation, serves no protective function ^2^. Under healthy conditions, the balance between excitation and inhibition is crucial for physiological pain. In the case of pathological pain, this balance is dysregulated, with enhanced excitation and reduced inhibition ^3, 4^.

An unappreciated tissue in the context of pathological pain is the meninges. Meninges, the three layers covering the brain and spinal cord ^5^, house a wide variety of immune cells, and these immune cells and their secreted proteins interact with cells of the parenchyma ^6^. Of the meningeal immune cells, meningeal macrophages are of the most well characterized cells in the central nervous system (CNS) meninges ^7, 8^. A subpopulation of meningeal macrophages has been identified that express the mannose receptor CD206 ^7, 8^. Whether these macrophages play a role in regulating basal nociception or hypersensitivity after injury remains elusive. In the present study, we focussed on the potential for meningeal CD206^+^ macrophages to be involved in pain processing by using a novel approach to target these cells in the meninges of male and female rats with and without skin injury.

## Materials and Methods

### Animals

Male and female Sprague-Dawley rats aged 6 to 12 weeks were obtained from Charles River Laboratories (Boucherville, Canada). Same-sex pairs of rats were housed in polycarbonate cages on a 14:10 hours light:dark cycle (lights on at 06:00 hours) in a temperature-controlled environment with *ad libitum* access to food and water. Experimenters were blinded, meaning that they were unaware of the nature of the substances that were being administered. Blinding to sex was not possible in behavioral experiments. At the end of behavioral experiments, animals were euthanized with inhaled CO_2_ and conformation of death was achieved by dislocation of the neck. For collecting tissue for bulk RNA sequencing and immunohistochemistry experiments, animals were deeply anaesthetized with urethane (2 g/kg, i.p.) and killed by decapitation.

### Ethical considerations

All experiments were approved by the Hospital for Sick Children’s Animal Care Committee and in compliance with the Canadian Council on Animal Care guidelines.

### Skin incision surgery

Rats were anaesthetized with 3% isoflurane/oxygen under sterile conditions. Using a scalpel no. 10, an incision was made on the left thigh, about one centimeter long, running parallel to, and immediately distal to the femur bone. The muscle underneath the skin was not touched, cut or in any way injured. The skin incision was closed using 6-0 vicryl sutures.

### Behaviour experiments

#### Controlled confounding factors for behaviour experiments

There are external factors that affect pain behaviour experiments, such as time of testing throughout the day, time of testing throughout the year, animal housing, the experimenter and their associated specifications, such as sex, scent and method of filament application. To control these factors, all our experiments were done at the same time in the day. Baseline (BL) experiments were always done from 9-10 am, and surgeries were done from 10 am to 12 pm. Post operative day 3 (POD3) measurements were always done from 9-10 am, and intrathecal injections were always done between 10-10:45 am. Four-hour time points were always around 2-2:45 pm, and 24-hour time points were always done between 9-10 am on the day after intrathecal injection. Therefore, for all experiments, for both sexes, timing of the day remained consistent. Further, the timing between each time point was exact, down to the minute. For example, between 1h timepoint and 2h timepoint, the experimenter waited 60 minutes, and between 2h timepoint and 4h timepoint, the wait time was 120 minutes for every animal being tested. MK was the experimenter for all pain behaviour measurements and skin incision surgeries. The animals were housed in the same room, under the same conditions, for the same amount of time, which was 1 week before experiments began. For experiments that tested both males and females, the sexes were tested separately, always one week apart. Male rats were tested first, and immediately the week after, female rats were tested. Animals were randomized in experimental groups and the behavioural experimenter was unaware as to treatment.

#### Von Frey testing

The mechanical withdrawal threshold of animals was tested on the ipsilateral and contralateral paw using calibrated von Frey filaments of increasing logarithmic nominal force values ^9^. Animals were placed in custom-made Plexiglas cubicles on a perforated metal floor and were permitted to habituate for 1 h before testing. Filaments were applied to the perpendicular plantar surface of the hind paw for 5 seconds. A positive response was recorded if there was a quick withdrawal, licking or shaking of the paw by the animal. Each filament was tested five times with increasing force filaments (1–15 g) used until a filament in which three out of five applications resulted in a paw withdrawal or when the maximal force filament was reached. This filament force is called the mechanical withdrawal threshold.

#### Hargreaves test

Thermal hyperalgesia was assessed by the Hargreaves test ^9^. In brief, animals were habituated in Plexiglas cubicles on a glass surface. The time (in seconds) for the hindpaw withdrawal from a radiant heat (IR = 45) stimulus projected to the plantar surface was measured. Three measurements on the ipsilateral hind paw were collected.

#### Acetone drop test

For cold assessment, 50 µL of acetone was applied to the midplantar surface of the hind paw and animal response was monitored for 20 seconds immediately after acetone application ^9^. The time (seconds) that the animal spent nursing in response to the cooling effect, including withdrawal, flick, stamp, or lick, was recorded. Three measurements were collected on the ipsilateral side, with each acetone application 5 minutes apart.

### Cell-depletion experiments

Cell depletion experiments were performed using biotinylated mannose-receptor (CD206) - antibody tagged to streptavidin. First, anti-rabbit mannose receptor antibody (Abcam, cat#ab64692) was biotinylated, following instructions of the Abcam biotinylation kit (ab201795). Briefly, 10 µl of modifying agent from the Abcam biotinylation kit was added to 100 µg (in 100 µl) of mannose receptor antibody. Modified antibody was then added to the biotin mix powder from the kit, and left at room temperature, in the dark, for 15 minutes to incubate. After 15 minutes, 10 µl of the quencher reagent was added to the mix, and the solution incubated for another 5 minutes at room temperature. Then, using streptavidin-ZAP kit (Advanced Targeting Systems IT-27), biotinylated mannose receptor antibody was tagged to saporin, which was converted into targeted toxin towards macrophages that express mannose receptor (mannose-receptor-saporin; referred to as CD206-saporin). Equal parts, in moles, streptavidin and mannose-receptor antibody were mixed (160 µl of biotinylated antibody mixed with 38 µl of streptavidin-ZAP 2.6 µg/µl). To control lot-to-lot variability and variation in freeze thaw, we used fresh biotinylation kit and antibody with each round of injections. Rats were injected intrathecally (30 µl) with saline, CD206-saporin (20 µg mannose-receptor antibody and 7µg of streptavidin-ZAP in 30 µl), or rabbit IgG conjugated to saporin. For the first cohort, rabbit IgG was used as the vehicle control. However, all experiments using rabbit IgG were repeated with phosphate-buffered saline (PBS) as vehicle control, unless otherwise stated in the figure caption.

### Drug administration

#### Intrathecal injection

Rats were anaesthetized with 3% isoflurane/oxygen under sterile conditions. The injection area was trimmed of all fur and wiped with 70% ethanol. All intrathecal injections were performed using a 50 µl Hamilton syringe with a 25-gauge beveled needle between the L5 and L6 vertebrae of anaesthetized rats.

#### Intraperitoneal injection

In a calm state, the tail of the rats was lifted gently until their lower abdomen was exposed. Using a 25-guage needle, the injection was performed on the lower abdomen into the intraperitoneal space. The rats remained calm throughout the procedure.

#### APV

D-(–)-2-Amino-5-phosphonopentanoic Acid (D-AP5, Sigma-Aldrich Cat#165304) was dissolved in PBS to a stock solution of 10 µg/µL. A fresh solution was prepared for intrathecal injection with a dose of 1.5 µg/10 µL. PBS was used as vehicle control.

#### CLP290

CLP was dissolved in 20% of 2-(Hydroxypropyl)-β-cyclodextrin (HPCD, Sigma-Aldrich Cat#H107) in PBS, and fresh solution was prepared daily. The dose for each animal was 100 mg/kg body weight of CLP290 (kindly donated by Yves De Konick, Université Laval). A solution of 20% of HPCD was used a vehicle control.

### Lumbar spinal cord meningeal dissection for immunohistochemistry

After perfusion with PBS followed by 4% paraformaldehyde, the vertebral column was separated from the dorsal spinal cord, starting at the cervical spinal cord and moving down to thoracic and lumbar spinal cord. Once the dorsal side of the lumbar spinal cord was exposed, the lumbar spinal cord was cut at the proximal and distal end and moved into a 35 mm dish with PBS. Under the microscope, dura and arachnoid were peeled off the lumbar spinal cord, and all peripheral nerves were separated to isolate the meninges. The meninges were then moved to an Eppendorf tube with PBS and stored, ready for immunohistochemistry.

### Immunohistochemistry

Male and female rats were terminally anaesthetized with urethane (1 ml of 20% urethane injected interperitoneally), and transcardially perfused with saline followed by 4% paraformaldehyde for immunohistochemical detection of microglia and meningeal macrophages. Meningeal macrophages were stained using anti-goat Iba1 (Cedarlane, cat#NB100-1028) and anti-rabbit mannose receptor 1 (anti-mannose receptor antibody from Abcam ab64692) antibodies. Parenchymal microglia were stained using Iba1 antibody. Cell counting of CD206^+^ macrophages was done by using ImageJ software.

### Lumbar dorsal root ganglion (DRG) 2-5 and lumbar spinal cord meningeal dissection for RNA extraction

Male and female rats were terminally anaesthetized with urethane (1 ml of 20% urethane injected interperitoneally), after which they were decapitated, and their blood was drained from the neck region under running water, to keep tissues fresh for RNA extraction. Then, the vertebral column was separated from the dorsal spinal cord, starting at the cervical spinal cord and moving down to thoracic and lumbar spinal cord. Once the dorsal side of the lumbar spinal cord was exposed, the lumbar spinal cord was cut at the proximal and distal end and moved into a 35 mm dish with RNALater. Under the microscope, dura and arachnoid were peeled off the lumbar spinal cord, and all peripheral nerves were separated to isolate the meninges. The meninges then are stored in an Eppendorf tube with RNALater, for future RNA extraction.

At the same time as meningeal dissection, DRG L2-L5 on the ipsilateral side of injured rats and the left side of naïve rats were located, extracted, and stored in an Eppendorf tube with RNALater, for future RNA extraction.

### RNA extraction for bulk RNA-sequencing

The Qiagen RNeasy kit was used to extract RNA from meninges and DRGs. Briefly, tissues were taken out of RNAlater and placed in new, clean Eppendorf tubes. Then, 150 µl of Qiazol was added to the Eppendorf tube, and the tissues sat in the Qiazol solution for 2 minutes, followed by tissue homogenization for 30-45 seconds. A final volume of 900 µl was reached with Qiazol and sat on the benchtop at room temperature for 5 minutes. Following, 100 µl of gDNA eliminator solution was added and the Eppendorf tube containing the tissue in solution was shaken vigorously by hand for 15 seconds. Chloroform (180 µl) was added and the Eppendorf tube containing the tissue in solution was shaken vigorously by hand for 15 seconds. The tube containing the homogenate sat on the benchtop at room temperature for 2-3 minutes. The mixture was centrifuged at 12,000 x *g* for 15 minutes at 4 degrees Celsius. The upper, aqueous phase was transferred to a new Eppendorf tube, and the same volume of 70% ethanol was added to the aqueous solution. The solution was then added to the spin columns (maximum 700 µl each round, for two rounds), and the column was centrifuged at room temperature for 15 seconds at 8000 x *g*. Flow-through was discarded for each round until the entire solution (ethanol and aqueous solution) went through the spin column. Buffer RWT (700 µl) was added to the spin easy column, the column was centrifuged at room temperature for 15 seconds at 8000 x *g*, and the flow-through was discarded. Buffer RPE (500 µl) was added to the spin easy column, the column was centrifuged at room temperature for 15 seconds at 8000 x *g*, and the flow-through was discarded. Another 500 µl of Buffer RPE was added to the spin easy column, the column was centrifuged at room temperature for 2 minutes at 8000 x *g*, and the flow-through was discarded. The spin column was added to a new Eppendorf tube, and centrifuged at full speed, at room temperature, for 5 minutes. The spin column was then moved to a clean collection tube, and 25 µl of RNase-free water was directly added to the spin column membrane. The spin column, with the water in the membrane, sat at room temperature for 7 minutes, and then was centrifuged at 8000 x *g* for 1 minute. Elution was repeated once to increase concentration of RNA extracted from meningeal tissue.

### Bulk RNA-sequencing Procedure

Animals were euthanized and DRG L2-3, DRG L4-5 and lumbar meninges (dura and arachnoid) were harvested 3 days post skin incision procedure to study transcriptional changes. RNA was extracted from the tissue and preserved in RNALater and the library was prepared and sequenced using IlluminaHiSeq 4000 by TCAG at The Hospital for Sick Children. The filtered reads were aligned to a reference genome using STAR ^10^. The genome used in this analysis was *Rattus norvegicus* assembly (Rnor_6.0) after quality control, it was calculated the log2(CPM) (counts-per-million reads) and ran principal component analysis. The differential gene expression analysis was done using edgeR ^11^ Bioconductor packages. Genes with adjusted P-value <0.01 and log2fold changes greater than |0.5| were defined as differentially expressed genes (DEGs). In this study a total of 66 samples were analysed. Six control groups (MNG_Naive_M, MNG_Naive_F, DRG23_Naive_M, DRG23_Naive_F, DRG45_Naive_M, DRG45_Naive_F) were used as a reference to find differentially expressed genes. Samples and pair-wise comparisons are summarized in Table 1 and Table 2, respectively.

**Table 1.** Summary of samples for RNA-sequencing experiment.

| **Sex** | **Condition** | **Tissue** |
| --- | --- | --- |
| Male (n=11) | Naïve (n=5) | Lumbar meninges |
|  |  | DRG L2-3 |
|  |  | DRG L4-5 |
|  | Skin incision injury (n=6) | Lumbar meninges |
|  |  | DRG L2-3 |
|  |  | DRG L4-5 |
| Female (n=11) | Naïve (n=5) | Lumbar meninges |
|  |  | DRG L2-3 |
|  |  | DRG L4-5 |
|  | Skin incision injury (n=6) | Lumbar meninges |
|  |  | DRG L2-3 |
|  |  | DRG L4-5 |

**Table 2.** Pair-wise comparisons for RNA-sequencing experiments.

| Pairwise comparisons - Males |  |  |
| --- | --- | --- |
| Naïve lumbar meninges | vs. | Skin incision lumbar meninges |
| Naïve DRG L2-L3 |  | Skin incision DRG L2-L3 |
| Naïve DRG L4-L5 |  | Skin incision DRG L4-L5 |
| Pair wise comparisons Females |  |  |
| Naïve lumbar meninges | vs. | Skin incision lumbar meninges |
| Naïve DRG L2-L3 |  | Skin incision DRG L2-L3 |
| Naïve DRG L4-L5 |  | Skin incision DRG L4-L5 |
| Pairwise comparisons – both sexes combined |  |  |
| Naïve lumbar meninges | vs. | Skin incision lumbar meninges |
| Naïve DRG L2-L3 |  | Skin incision DRG L2-L3 |
| Naïve DRG L4-L5 |  | Skin incision DRG L4-L5 |

### Statistical analysis

Data are presented as means ± SEM. Statistical analysis was performed using GraphPad Prism 9, with statistical significance set as p < 0.05. Statistical tests are described in figure legends for each corresponding experiment.

## Results

### Depleting CD206^+^-meningeal macrophages in spinal cord lumbar meninges via intrathecal administration of CD206-saporin

We investigated the necessity of CD206^+^ macrophages in the lumbar meninges by an approach to deplete these cells using intrathecally administered anti-CD206 conjugated to the toxin saporin (subsequently called ‘CD206-saporin’). To determine whether CD206-saporin intrathecal injection causes depletion of meningeal CD206^+^ macrophages, we harvested lumbar meningeal tissue (dura and arachnoid) from male and female rats with either intrathecal CD206-saporin injection or intrathecal PBS injection as control (Figure 1A). We found that the number of CD206^+^ cells in the lumbar meninges of CD206-saporin-injected rats was significantly less than that of PBS-injected rats (Figure 1B and 1C). Thus, we conclude that meningeal CD206^+^macrophages were depleted in male and female rats after CD206-saporin intrathecal injection.

**Figure 1.**
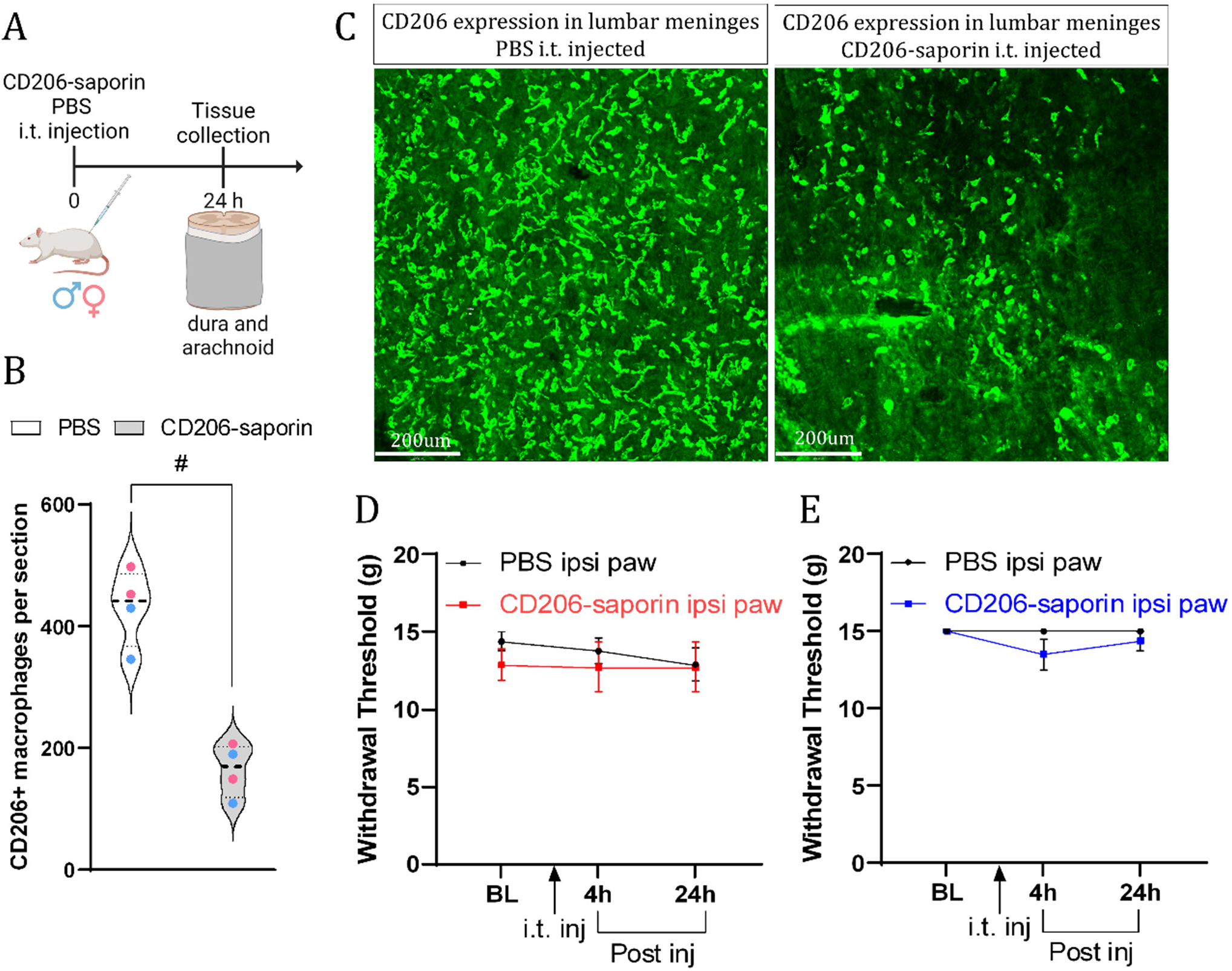
Intrathecal administration of CD206-saporin, and thus depletion of CD206^+^ meningeal macrophages, does not affect mechanical sensitivity in naïve rats. (A) Experimental outline. (B) Quantification of CD206^+^ macrophages in lumbar dura and arachnoid 1 day post injection. n = 4 (2M 2F) PBS and 4 (2M 2F) CD206-saporin injected animals, respectively. # signifies P-value < 0.05, two-sided Student’s t-tests. Pink dots signify female rats and blue dots signify male rats. (C) Representative images of meningeal macrophages expressing CD206, one day post-injection of PBS, CD206-saporin. Scale bar = 200 μm. i.t.: intrathecal. (D) Line graph of mechanical withdrawal threshold in naïve female rats at baseline and 4h and 24h after intrathecal injection of PBS and CD206-saporin. n=8 female rats per condition. (E) Line graph of mechanical withdrawal threshold in naïve male rats at baseline and 4h and 24h after intrathecal injection of PBS and CD206-saporin. n=7 and 8 from CD206-saporin and PBS treated male rats, respectively. Data presented as mean ± SEM. Mann Whitney U test comparing PWT at each time point between CD206-saporin-and PBS-injected groups. BL: baseline, i.t. inj: intrathecal injection, inj: injection.

### Intrathecal administration of CD206-saporin does not alter mechanical sensitivity in naïve male or female rats

Before exploring whether depleting CD206^+^ macrophages in the meninges affects pain sensitivity after injury, we investigated whether these cells influence basal nociception in naïve female and male rats by testing their sensitivity to innocuous mechanical stimuli ^12^. Following CD206-saporin administration, we assessed sensitivity to mechanical stimuli of the hind paw of naïve male and female rats, using Von Frey filaments of varying bending force and measured their paw withdrawal threshold (PWT) 4h after macrophage depletion and then again 24h later. For the control experiment, male and female rats received intrathecal injection of PBS.

In females, there was no statistically significant difference between PWT of rats injected with PBS compared to PWT of rats injected with CD206-saporin at baseline (BL, before intrathecal injection) (Figure 1D). Similarly, there was no significant different between PWT of female rats injected with PBS compared to female rats injected with CD206-saporin 4h and 24h after injection. In males, we found that there was no statistically significant difference between baseline PWT (BL, before intrathecal injection) of rats injected with PBS compared to PWT of rats injected with CD206-saporin at (Figure 1E). In addition, there was no significant difference between PWT of male rats injected with PBS compared to male rats injected with CD206-saporin 4h and 24h after injection. Taking these findings together we conclude that depleting meningeal macrophages by intrathecal injection of CD206-saporin does not alter mechanical sensitivity in naïve rats of either sex.

### Intrathecal administration of CD206-saporin after skin incision causes mechanical sensitization in male but not female rats

Having determined that depleting CD206^+^ macrophages in the meninges does not impact basal mechanical sensitivity in naïve male and female rats, we proceeded to examine the potential role of these macrophages in sensitivity to mechanical stimuli after minor skin incision injury on the upper left thigh, which is the injury typically used as the ‘sham’ in studies of peripheral nerve injury ^13–15^. First, we measured PWT of male and female rats before and 3 days after skin incision injury (post operative day 3, POD3). There was no statistically significant difference between PWT of male and female rats at POD3 compared to their PWT at baseline. Thus, we found that minor skin incision injury on the upper left thigh did not affect PWT, and therefore mechanical sensitivity, in the ipsilateral hind paw of male and female rats (Figure 2).

**Figure 2.**
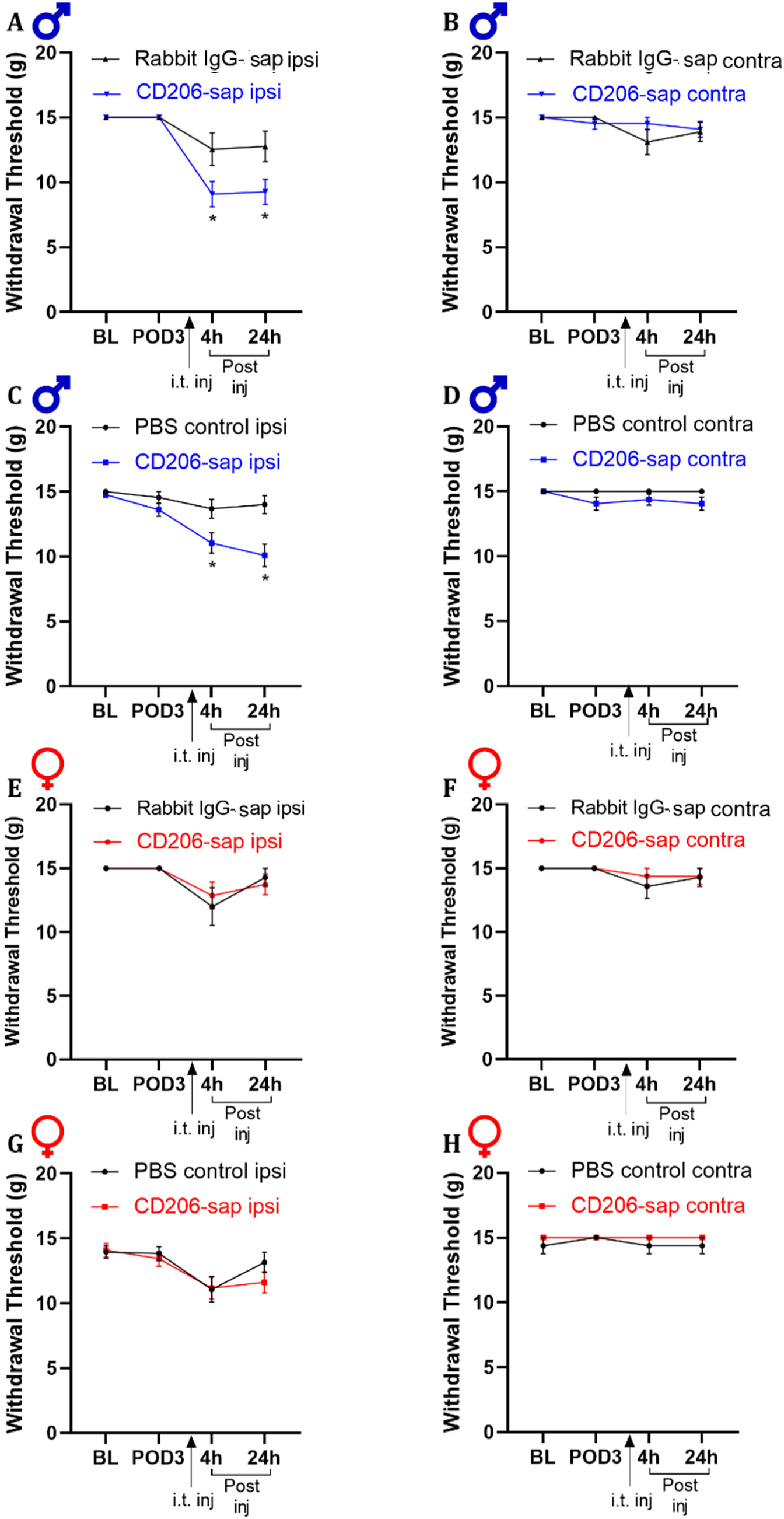
Intrathecal administration of CD206-saporin causes mechanical sensitivity in male, but not female rats after skin incision. (A) Line graph of mechanical withdrawal threshold in the ipsilateral paw of male rats at baseline (BL), 3days after skin incision surgery (POD3) and 4h and 24h after intrathecal injection of rabbit IgG-saporin and CD206-saporin. n=11 and 9 from CD206-saporin and rabbit-IgG treated male rats, respectively. (B) Line graph of mechanical withdrawal threshold in the contralateral paw of male rats at baseline (BL), 3days after skin incision surgery (POD3) and 4h and 24h after intrathecal injection of rabbit IgG-saporin and CD206-saporin. n=11 and 9 from CD206-saporin and rabbit-IgG treated male rats, respectively. (C) Line graph of mechanical withdrawal threshold in the ipsilateral paw of male rats at baseline (BL), 3days after skin incision surgery (POD3) and 4h and 24h after intrathecal injection of PBS and CD206-saporin. n=23 and 16 from CD206-saporin and PBS treated male rats, respectively. (D) Line graph of mechanical withdrawal threshold in the contralateral paw of male rats at baseline (BL), 3days after skin incision surgery (POD3) and 4h and 24h after intrathecal injection of PBS and CD206-saporin. n=16 and 8 from CD206-saporin and PBS treated male rats, respectively. (E) Line graph of mechanical withdrawal threshold in the ipsilateral paw of female rats at baseline (BL), 3days after skin incision surgery (POD3) and 4h and 24h after intrathecal injection of rabbit IgG-saporin and CD206-saporin. n=7 and 8 from CD206-saporin and rabbit-IgG treated male rats, respectively. (F) Line graph of mechanical withdrawal threshold in the contralateral paw of female rats at baseline (BL), 3days after skin incision surgery (POD3) and 4h and 24h after intrathecal injection of rabbit IgG-saporin and CD206-saporin. n=7 and 8 from CD206-saporin and rabbit-IgG treated male rats, respectively. (G) Line graph of mechanical withdrawal threshold in the ipsilateral paw of female rats at baseline (BL), 3days after skin incision surgery (POD3) and 4h and 24h after intrathecal injection of PBS and CD206-saporin. n=23 and 19 from CD206-saporin and PBS treated female rats, respectively. (H) Line graph of mechanical withdrawal threshold in the contralateral paw of female rats at baseline (BL), 3days after skin incision surgery (POD3) and 4h and 24h after intrathecal injection of PBS and CD206-saporin. n=8 and 16 from CD206-saporin and PBS treated female rats, respectively. Data presented as mean ± SEM. Mann Whitney U test comparing PWT at each time point between CD206-saporin- and PBS-injected groups. * = P <0.05. BL: baseline, POD: post-operative day, i.t. inj: intrathecal injection.

After measuring PWT on POD3, CD206-saporin or rabbit IgG-saporin was intrathecally administered. Because macrophages and microglia are phagocytic cells ^8^, a concern we had with CD206-saporin usage was that all macrophages and microglia were phagocytosing saporin, independent of CD206 antibody. To address this question, we used a non-specific rabbit IgG tagged to saporin, and determined whether injection of saporin tagged to a rabbit IgG has the same effect as CD206-saporin. As shown in Figure 2A, following CD206-saporin there was a significant decline in PWT in male rats 4h and 24h post-injection. However, the rabbit IgG-saporin control did not cause a significant reduction in PWT after administration compared to before administration. Thus, administrating CD206-saporin, but not rabbit IgG-saporin causes mechanical hypersensitivity in male rats with skin incision.

As a second control in male rats with skin incision injury, we compared PWT after intrathecal injection of CD206-saporin to intrathecal injection of PBS. We found that in males with skin incision injury, PWT of those intrathecally injected with CD206-saporin was significantly lower than PWT of those intrathecally injected with PBS, 4h and 24h post-depletion (Figure 2C). Thus, intrathecal injection of CD206-saporin caused a significant decline in PWT of male rats with skin incision injury, while intrathecal injection of PBS did not affect PWT in male rats with skin incision injury. Taking these findings together, we conclude that after skin incision injury, intrathecal injection of CD206-saporin in the meninges unmasks mechanical sensitivity 4h and 24h after injection in male rats.

By contrast, in female rats with skin incision injury intrathecal injection transiently decreased PWT of both CD206-saporin-injected and rabbit IgG-saporin-injected group (Figure 2E). However, there was no statistically significant difference in PWT of female rats injected with rabbit IgG-saporin compared to female rats injected with CD206-saporin either at 4h or at 24h after administration. These findings are taken as evidence that in female rats with skin incision injury, intrathecal injection of CD206-saporin does not cause mechanical sensitivity 4h or 24h after administration.

Moreover, in a separate cohort of female rats with skin incision injury, intrathecal injection transiently decreased PWT in both CD206-saporin-injected group and in the second control group – PBS-injected (Figure 2G). There was no statistically significant difference in PWT of female rats injected with PBS compared to female rats injected with CD206-saporin 4h or 24h after administration. Thus, in contrast to our findings in male rats with skin incision, in female rats with skin incision injury, intrathecal injection of CD206-saporin does not affect mechanical sensitivity 4h and 24h after administration.

### Intrathecal administration of CD206-saporin does not affect widespread mechanical sensitivity in male and female rats after skin incision

Thus far, we have determined that administration of CD206-saporin results in mechanical sensitivity in the ipsilateral paw in male rats with skin incision injury 4h and 24h after CD206-saporin administration. To investigate whether intrathecal administration of CD206-saporin may have a widespread effect on mechanical sensitivity in animals with skin incision injury, we measured PWT in the contralateral paw.

In male and female rats with skin incision injury, we found that PWT was not different after CD206-saporin administration compared to after rabbit IgG-saporin or after PBS administration (Figure 2B, D, F, H). Thus, administering CD206-saporin did not affect PWT in the contralateral paw in male or female rats with skin incision injury, suggesting that there is no widespread effect on mechanical sensitivity of injecting CD206-saporin in male or female rats with skin incision injury.

### Intrathecal administration of CD206-saporin does not affect heat or cold sensitivity in male or female rats after skin incision

In addition to sensitivity to mechanical stimuli, another indicator of pain is sensitivity to innocuous thermal stimuli (thermal allodynia) or heightened sensitivity to noxious thermal stimuli ^9^. Thus far, we have found that intrathecal administration of CD206-saporin in the meninges leads to sensitization to mechanical stimulation in male rats. In the following experiments we evaluated sensitivity to cold or heat stimuli in male and female rats with skin incision injury after CD206-saporin intrathecal administration.

To characterize the effects of CD206-saporin intrathecal administration on cold sensitivity in male and female rats with skin incision injury, we measured response duration after the hind paw of the rats was exposed to a drop of acetone (50 µl) (Figure 3A and B). At baseline, none of the male rats responded to acetone exposure, while females showed limited response. After skin incision injury (POD3), there were no changes compared to baseline in male or female rats. Thus, the skin incision injury alone did not cause cold sensitivity in male or female rats.

**Figure 3.**
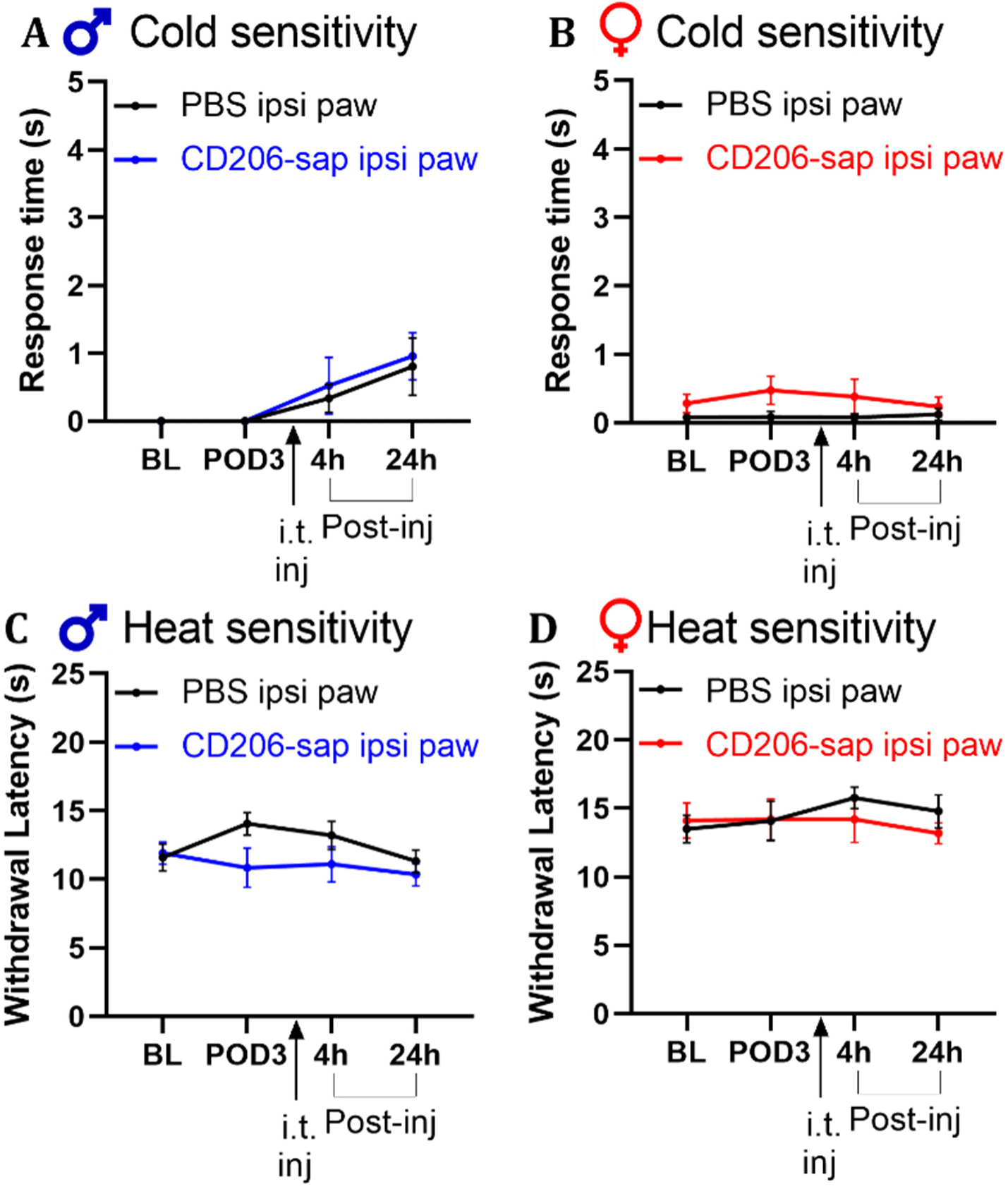
Male and female rats with skin incision are not sensitive to thermal stimuli after CD206-saporin intrathecal injection on POD3. (A) Cold sensitivity in male rats at baseline (BL), 3days after skin incision surgery (POD3) and 4h and 24h after intrathecal injection of PBS and CD206-saporin. n=7 and 8 from CD206-saporin and PBS treated male rats, respectively. (B) Cold sensitivity in female rats at baseline (BL), 3days after skin incision surgery (POD3) and 4h and 24h after intrathecal injection of PBS and CD206-saporin. n=7 and 8 from CD206-saporin and PBS treated female rats, respectively. (C) Heat sensitivity in male rats at baseline (BL), 3days after skin incision surgery (POD3) and 4h and 24h after intrathecal injection of PBS and CD206-saporin. n=7 and 8 from CD206-saporin and PBS treated male rats, respectively. (D) Heat sensitivity in female rats at baseline (BL), 3days after skin incision surgery (POD3) and 4h and 24h after intrathecal injection of PBS and CD206-saporin. n=7 and 8 from CD206-saporin and PBS treated female rats, respectively. Data presented as mean ± SEM. Mann Whitney U test comparing PWT at each time point between CD206-saporin- and PBS-injected groups. * = P <0.05. BL: baseline, POD: post-operative day, i.t. inj: intrathecal injection.

After intrathecal injection we quantified cold responses in the males (Figure 3A). There were no statistically significant differences in the animals that had received intrathecal CD206-saporin as compared with those that had received intrathecal PBS at either 4h or 24h timepoint (Figure 3A). Likewise, in females we found no differences in cold responses in CD206-saporin-treated versus PBS-treated animals (Figure 3B). From these findings, we conclude that intrathecal administering CD206-saporin after skin incision injury does not induce cold sensitization in either sex.

We evaluated heat sensitivity in each sex by measuring withdrawal latency where the hind paws of rats were exposed to infrared light, before and after skin incision injury and after CD206-saporin and PBS intrathecal administration (Figure 3C and D). The baseline response latencies in animals that went on the receive intrathecal CD206-saporin were not different from those that went on to receive PBS males or in females. Moreover, on POD3, withdrawal latency was not significantly different compared to baseline in male or female rats. Thus, skin incision alone did not affect withdrawal latency in either sex.

Furthermore, we found in male and in female rats with skin incision injury, there was no significant difference in withdrawal latency between the group with that received intrathecal CD206-saporin versus the control group with intrathecal PBS at either 4h or 24h after administration (Figure 3C and D). Thus, we conclude that CD206-saporin intrathecal administration did not affect withdrawal latency in male or female rats with skin incision injury.

### Blocking NMDARs reverses mechanical sensitivity induced by intrathecal administration of CD206-saporin in male rats with skin incision

Thus far, we have found in male, but not female, rats with skin incision injury that intrathecal administration of CD206-saporin selectively unveils sensitization to mechanical but not to thermal stimulation. Therefore, we focused on determining spinal mechanisms that may underlie this male-only mechanical sensitization. As pathological pain is characterized in the spinal cord by dysregulation of the balance between excitation, which is enhanced, and inhibition which is reduced ^3, 4^, we investigated the role of each of these induced by intrathecal administration of CD206-saporin after skin incision in male rats.

Increased activity of NMDA receptors (NMDARs) in the spinal dorsal horn has been shown to mediate pain hypersensitivity in many models of chronic pain ^2, 16^. To investigate spinal NMDARs in the mechanical sensitization induced by CD206-saporin we intrathecally administered the NMDAR competitive antagonist 2-amino-5-phosphonovaleric acid (APV), or PBS vehicle control, 24 hours after the animals had received CD206-saporin (Figure 4A). At this timepoint, when the mechanical hypersensitivity was robustly established, we found that intrathecally administered APV increased the PWT that had been reduced by CD206-saporin (Figure 4A). Therefore, we conclude that APV reversed the sensitization, and thus that NMDARs are necessary for the CD206-saporin-induced mechanical sensitization in male rats with skin incision.

**Figure 4.**
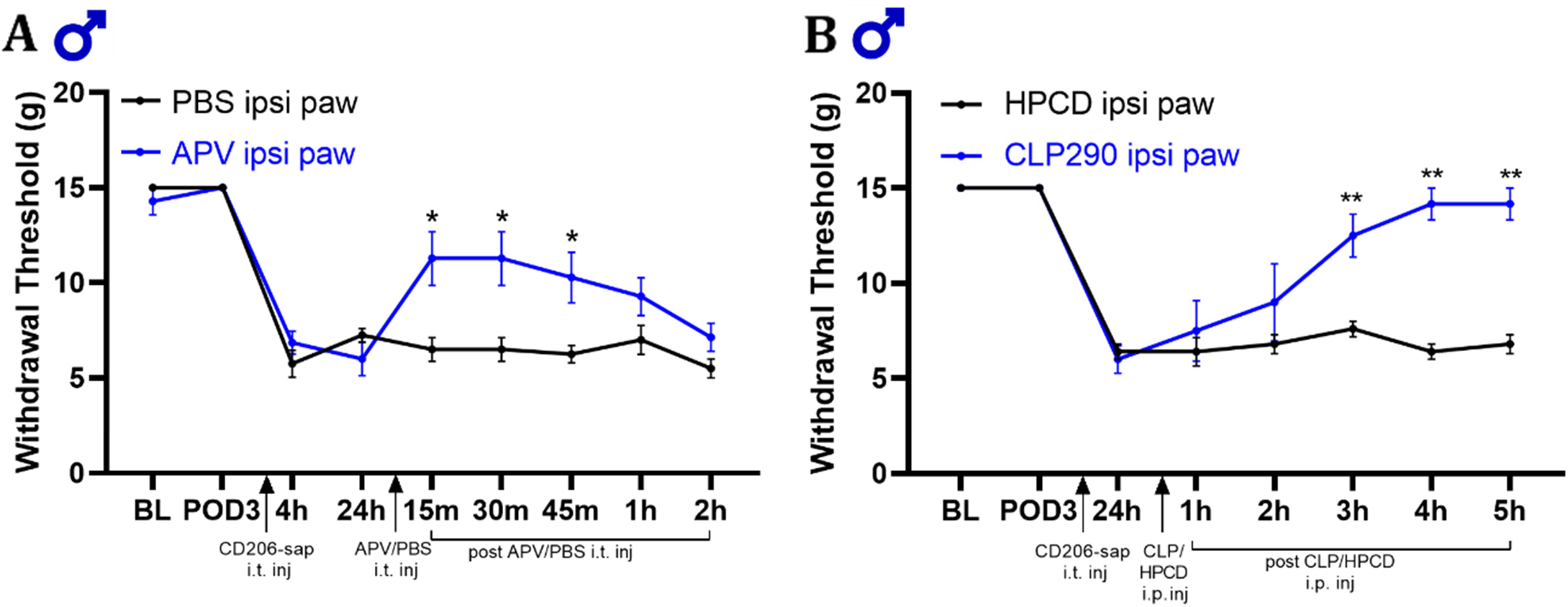
Blocking NMDAR or activating KCC2 reverses mechanical sensitization in male rats with skin incision injury after CD206-saporin intrathecal administration. (A) Line graph of mechanical paw withdrawal in male rats at baseline (BL), days after skin incision procedure (POD3), 4 and 24 hours after meningeal CD206+-macrophage depletion, and 15 mins, 30 mins, 45 mins, 1h and 2h after intrathecal injection of NMDA receptor blocker APV or PBS. n = 7 for both groups. (B) Line graph of mechanical sensitivity in male rats at baseline (BL), 3 or 4 days after skin incision procedure (POD3-4), 24 h after meningeal CD206+-macrophage depletion, and 1, 2, 3, 4, and 5 h intrathecal injection of KCC2 activator CLP290 (100 mg/kg) or 20% HPCD. n = 5 for CLP290 group and n = 6 for PBS group. Data presented as mean ± SEM. Mann Whitney U test comparing PBS to APV group or HPCD to CLP290 group at each timepoint. ** = P <0.01, * = P <0.05. BL: baseline, POD: post-operative day, i.t. inj: intrathecal injection, i.p inj: intraperitoneal injection, HPCD: 20% 2-hydroxypropyl- β-cyclodextrin.

### Activating potassium-chloride co-transporter (KCC2) reverses mechanical sensitivity induced by intrathecal administration of CD206-saporin in male rats with skin incision

Peripheral nerve damage is associated with reduced KCC2 activity in the lamina I spinal cord dorsal horn neurons, which causes a depolarizing shift in the anion reversal potential by increasing intracellular Cl^-^, thereby diminishing synaptic inhibition ^17^. Increasing KCC2 function pharmacologically decreases raised intracellular Cl^-^ concentration, thereby rescuing synaptic disinhibition resulting in reversal of pain hypersensitivity ^18, 19^. Therefore, we investigated the possibility that disinhibition by suppressing KCC2 function may also be necessary for the mechanical sensitivity induced by intrathecal administration of CD206-saporin after skin incision in male rats. For this purpose, we tested the molecule CLP290, a prodrug for the active metabolite CLP257 ^18^,which after intraperitoneal administration has been shown to reversal pathological pain hypersensitivity in rats ^19^ (Figure 4B). Three days after skin incision injury (POD3) CD206-saporin was intrathecally injected in male rats. Twenty-four hours later, when the mechanical sensitization was robust, we administered CLP290 (i.p.), or its vehicle control. We found that intraperitoneal injection of CLP290, but not the vehicle control, increased the PWT that had been lowered by administering CD206-saporin (Figure 4B). Therefore, we conclude that activating KCC2 with CLP290 reverses mechanical sensitization induced by CD206-saporin administration in male rats with skin incision injury.

### Transcriptional analysis of lumbar meninges and DRGs in rats with or without skin incision injury

From our findings that in male rats intrathecal administration of CD206-saporin unveils mechanical sensitization after skin incision but not in naïve animals, and that CD206-saporin depletes CD206^+^ macrophages, we infer that the skin incision in the thigh induces sensitization to mechanical stimuli applied to the paw but that this sensitization is masked by a response induced in these cells in the meninges. The input from the periphery to drive changes in the spinal cord comes via the primary afferents innervating the skin where the incision is made, the cells bodies of which in lumbar dorsal root ganglia (DRG) L2-L3 ^20^. On the other hand, the stimuli that drive the behavioural responses are applied to the hind paw, which is innervated by primary afferents having their cell bodies in DRG L4-L5. From these considerations together, it is conceivable that the masking of the mechanical sensitization may involve molecular changes in cells in the meninges, in DRG2-3, or in DRG4-5. We wondered therefore whether we could uncover the locus, or loci, of relevant changes by alterations in the transcriptional programing in these three tissues. To explore this idea, we interrogated the transcriptomes of lumbar meninges, DRG2-3, and DRG4-5 using bulk RNA sequencing (see Figure 5A).

**Figure 5.**
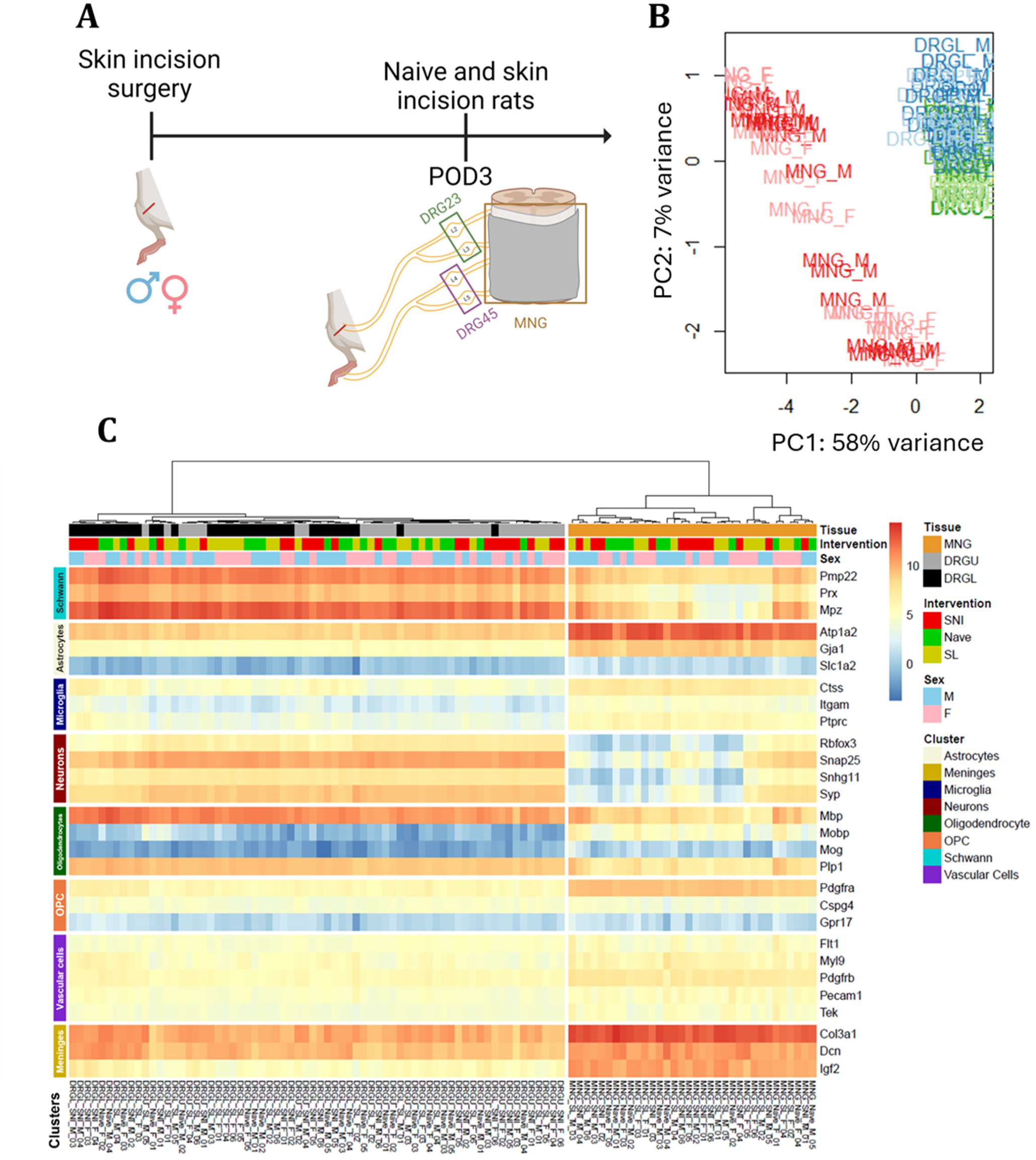
Principal component analysis (PCA) shows clear clustering of meninges (MNG) and DRGs. (A) Experimental design and overview. (B) Principal component analysis in tissues and sex. Scatter plot representing the scores along the first principal component (PC1) and the second component principal (PC2). (C) Heatmap showing the expression of genes that are specific to different tissue and cell types surrounding the spinal cord. POD: post-operative day, FC: fold change, dim: dimension, MNG = meninges, DRGU = DRG L2-L3, DRGL = DRG L4-L5, F = female, M = male.

To define main sources of transcriptome variability, we first analyzed the datasets at the sample level by principal component analysis (PCA) (Figure 5B). The principal components 1 (58%) and 2 (7%) were the major principal components, accounting for 65% of the overall variance. We found that there was a clear clustering of the samples from meninges (MNG) and DRG. Interestingly, no sex-dependent clustering of samples from male or female rats was observed at the level of the PCA (Figure 5B).

Before investigating differentially expressed genes, we assessed the quality of the samples ^21^ to ensure that they were not contaminated with tissue from the spinal cord parenchyma (Figure 5C). We found that meningeal tissue was enriched with transcripts specific to the meninges, as well as *Atp1a2*. Although *Atp1a2* is highly expressed in astrocytes ^22^ and typically used as a marker for astrocytes, it is also expressed in other types of tissue, including vascular tissue ^23^. Since meningeal tissue is not enriched with *Slc1a2* (Figure 5C) another transcript specific to astrocytes ^24^, we conclude that there were no astrocytes in the sample. Further, transcripts specific to neurons and oligodendrocytes had very low or no expression in meningeal tissue. Therefore, there were no evidence of substantive contamination by spinal cord parenchymal tissue in the samples of lumbar meninges. Likewise, the DRG samples had very low expression of transcripts specific to microglia and astrocytes (Figure 5C). Therefore, we conclude that there was no detectable spinal cord parenchymal tissue contaminant in the samples of the DRGs.

### Minimal effect of skin incision on gene expression in DRGs of rats of either sex

To determine whether skin incision alters gene expression in the DRGs and in the meninges we compared, by sex, transcriptomes of these tissues from naïve rats with those from rats three days after the skin incision (Figure 6). Differentially expressed genes (DEGs) were defined as the genes that pass the criteria of (EdgeR) adjusted p-value<0.05 and log2(FC) >|0.5|. With the DRG L2-3 transcriptomes, we found no DEGs when comparing male rats with skin incision to naïve male rats (Figure 6D). Nor did we find any DEGs when comparing female rats with skin incision to naïve female rats (Figure 6E). However, when the data from both sexes were combined, there were 8 genes that were higher in mRNA expression in the DRG L2 and L3 of rats with skin incision injury relative to naïve rats (Figure 6F, Supplementary Table 2). Similarly, no DEGs were found in DRGs L4-L5 in either sex comparing rats with skin incision to naïve (Figures 6G, H and I). From these findings we conclude that in males and in females three days after skin incision surgery, there were no detectable alterations in the transcriptomes either in DRG L2-L3, where the cell bodies of primary afferents that innervate the site of injury reside, or in DRG L4-L5 where the cell bodies of primary afferents that innervate the hind paw reside.

**Figure 6.**
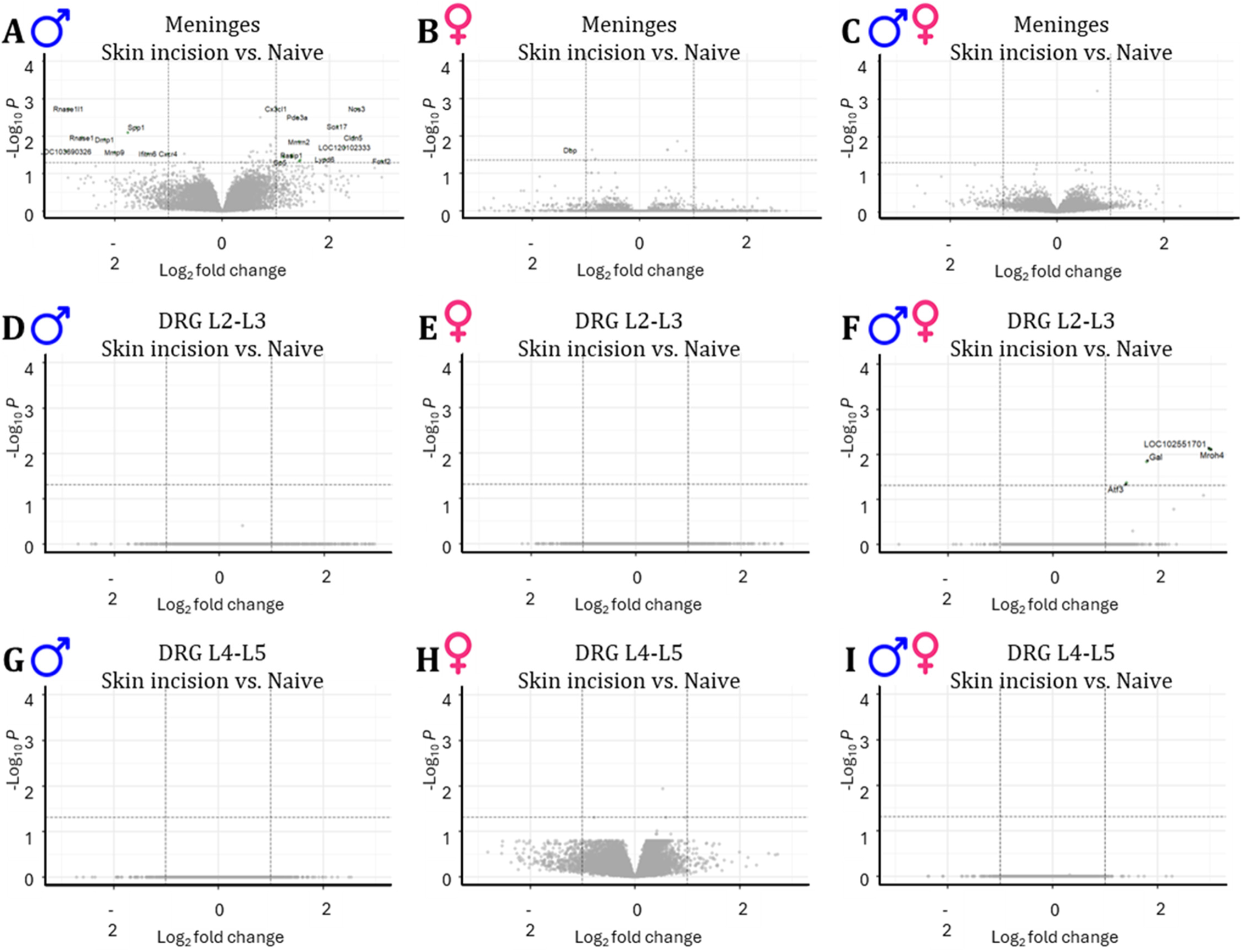
After skin incision injury, there are transcriptional changes in the male, but not female, lumbar meninges. (A) Volcano plot showing differentially expressed genes in lumbar meninges of male rats with skin incision vs naïve male rats. Volcano plots were obtained by plotting the log2 fold change (FC) of MNG_M.SL against the negative log10 of the EdgeR adjusted P-value. Genes that changed 1 log2(FC) or more with a significant of adjusted P-value <0.05 are shown in green. n = 6 for skin incision male rats and n = 5 for naïve male rats. (B) Volcano plot showing differentially expressed genes in lumbar meninges of female rats with skin incision vs naïve female rats. Volcano plots were obtained by plotting the log2 FC of MNG_F.SL against the negative log10 of the EdgeR adjusted P-value. Genes that changed 1 log2(FC) or more with a significant of adjusted P-value <0.05 are shown in green. n = 6 for skin incision female rats and n = 5 for naïve female rats. (C) Volcano plot showing differentially expressed genes in lumbar meninges of male and female rats with skin incision combined vs naïve male and female rats combined. Volcano plots were obtained by plotting the log2 FC of MNG_SL against the negative log10 of the EdgeR adjusted P-value. Genes that changed 1 log2(FC) or more with a significant of adjusted P-value <0.05 are shown in green. n = 6 for skin incision female rats and n = 5 for naïve female rats. (D) Volcano plot showing differentially expressed genes in DRG L2-L3 of male rats with skin incision vs naïve male rats. Volcano plots were obtained by plotting the log2 fold change (FC) of DRGU_M.SL against the negative log10 of the EdgeR adjusted P-value. Genes that changed 1 log2(FC) or more with a significant of adjusted P-value <0.05 are shown in green. n = 6 for skin incision male rats and n = 5 for naïve male rats. (E) Volcano plot showing differentially expressed genes in DRG L2-L3 of female rats with skin incision vs naïve female rats. Volcano plots were obtained by plotting the log2 FC of DRGU_M.SL against the negative log10 of the EdgeR adjusted P-value. Genes that changed 1 log2(FC) or more with a significant of adjusted P-value <0.05 are shown in green. n = 6 for skin incision female rats and n = 5 for naïve female rats. (F) Volcano plot showing differentially expressed genes in DRG L2-L3 of male and female rats with skin incision combined vs naïve male and female rats combined. Volcano plots were obtained by plotting the log2 FC of DRGU_M.SL against the negative log10 of the EdgeR adjusted P-value. Genes that changed 1 log2(FC) or more with a significant of adjusted P-value <0.05 are shown in green. n = 6 for skin incision female rats and n = 5 for naïve female rats. (G) Volcano plot showing differentially expressed genes in DRG L4-L5 of male rats with skin incision vs naïve male rats. Volcano plots were obtained by plotting the log2 fold change (FC) of DRGL_M.SL against the negative log10 of the EdgeR adjusted P-value. Genes that changed 1 log2(FC) or more with a significant of adjusted P-value <0.05 are shown in green. n = 6 for skin incision male rats and n = 5 for naïve male rats. (H) Volcano plot showing differentially expressed genes in DRG L4-L5 of female rats with skin incision vs naïve female rats. Volcano plots were obtained by plotting the log2 FC of DRGL_M.SL against the negative log10 of the EdgeR adjusted P-value. Genes that changed 1 log2(FC) or more with a significant of adjusted P-value <0.05 are shown in green. n = 6 for skin incision female rats and n = 5 for naïve female rats. (I) Volcano plot showing differentially expressed genes in DRG L4-L5 of male and female rats with skin incision combined vs naïve male and female rats combined. Volcano plots were obtained by plotting the log2 FC of DRGL_M.SL against the negative log10 of the EdgeR adjusted P-value. Genes that changed 1 log2(FC) or more with a significant of adjusted P-value <0.05 are shown in green. n = 6 for skin incision female rats and n = 5 for naïve female rats.

### Skin incision induces changes expression of numerous genes in the meninges of male, but not female rats

In contrast to the DRGs, we discovered alterations in the transcriptomes of the lumbar meninges three days after skin incision (Figure 6A, B and C). With meninges from male rats, we identified fourteen genes that were highly expressed with skin incision relative to naïve (Figure 6A and Supplementary Table 1). Ten genes were identified that were lower in gene expression in males with skin incision injury compared to naïve male rats (Figure 6A). In females, there was one gene that was lower in expression in rats with skin incision injury compared to naïve rats (Figure 6B). Taking these findings together, we conclude that there is a sex difference in the effect of skin incision on the lumbar meningeal transcriptomes, with a much larger and more diverse effect in males than in females. The differential effect on the meningeal transcriptome in males coincides with the masking of mechanical hypersensitivity in the paw three days after skin incision to the thigh.

## Discussion

Here, we discovered a role for CD206 macrophages in regulating mechanical hypersensitivity in male, but not in female, rats with skin incision injury. Administering CD206-saporin unmasked mechanical hypersensitivity in males but not females. In as much as CD206-saporin depleted CD206-expressing macrophages, the most parsimonious interpretation is that skin incision in the thigh produces hypersensitivity in the ipsilateral paw that is revealed by depleting CD206^+^ meningeal macrophages. We also found that intrathecal administration of CD206-saporin, and therefore depletion of CD206^+^ meningeal macrophages, does not affect mechanical sensitivity in the contralateral paw. Therefore, depletion of these macrophages does not have a widespread effect on mechanical sensitivity in male or female rats with skin incision injury. Intriguingly, the depletion of CD206^+^ macrophages in the meninges does not alter sensitivity to cold and heat stimuli, highlighting a specificity in their role towards mechanical pain hypersensitivity modulation.

In the healthy spinal cord dorsal horn, a delicate balance between the excitatory and inhibitory inputs ensures proper functioning of the somatosensory system, and dysregulated excitatory and inhibitory inputs contribute to the development of pain hypersensitivity ^4, 16, 25, 26^. Meningeal CD206^+^ macrophages may mask hypersensitivity through maintaining homeostasis between inhibitory and excitatory inputs in the spinal cord dorsal horn after skin incision injury. In this study, we determined that mechanical pain hypersensitivity induced by CD206-saporin intrathecal injection in injured male rats is reversed by both inhibition of NMDARs and activation of KCC2. Thus, depleting CD206^+^ meningeal macrophages unmasks hyperexcitation in the spinal cord due to increased NMDAR activity and decreased KCC2 activity. Taking these findings together, we suggest that meningeal CD206^+^ macrophages maintain spinal cord synaptic homeostasis after injury, by inhibiting NMDAR activity and activating KCC2.

Our findings here are evidence that meningeal CD206^+^ macrophages silence mechanical hypersensitivity after minor skin injury through action in the spinal cord. These results compliment studies in which it has been determined that peripheral macrophages expressing the CD206 receptor suppress mechanical hypersensitivity through actions at the site of injury ^27–29^. These peripheral anti-inflammatory macrophages have been shown to secrete opioid peptides Met-enkephalin, dynorphin A1-17 and β-endorphins when adoptively transferred at the site of nerve injury, and thereby reducing mechanical hypersensitivity ^27–29^. CD206^+^ macrophages may also shuttle mitochondria to sensory neurons, and therefore facilitate pain resolution ^30^. In the meninges, macrophages may exert their pro-resolution effects through enhancing IL10 production^31^. Thus, it is possible that through one of these mechanisms meningeal CD206^+^ macrophages may contribute to maintaining excitatory and inhibitory balance which masks mechanical hypersensitivity after skin injury.

Our findings also indicate that there are modality-specific effects of CD206^+^ meningeal macrophage depletion. In the periphery, macrophages in the DRG drive mechanical rather than cold allodynia ^32^, consistent with the observation that activated macrophages are primarily found around injured large-dimeter A-fiber sensory neuron cell bodies after sciatic nerve injury ^33^. Peripheral CD206^+^ macrophages, adoptively transferred to the injured sciatic nerve, reverse mechanical hypersensitivity, but did not affect heat hypersensitivity ^29^. These findings indicate that macrophages only affect mechanical hypersensitivity after nerve injury, complementing our observations that meningeal CD206^+^ macrophages only play a role in silencing mechanical hypersensitivity.

To date, no studies have identified sex-dependent mechanisms underlying the pro-resolution properties of anti-inflammatory macrophages in pain hypersensitivittable ^28, 34^. However, these studies have only used male rodents ^27, 29, 31^. Thus, the current study is the first to investigate the effects of CD206^+^ macrophages in the meninges of females with or without injury. When considering the sex-dependent contributions of macrophages in pain, work in the periphery has found that pain hypersensitivity is increased by distinct pathways in males and females. ^35^ ^36^. Sexual dimorphism has also been shown in the spinal dorsal horn where microglia are not involved in the development of nerve injury-induced pain hypersensitivity in females, whereas they contribute to the development of neuropathic pain in males ^37^. Instead, CD8^+^ T cells are involved in the development of neuropathic pain in females ^38^. Taking together, our findings show that meningeal macrophages exert an analgesic effect after skin incision injury in a sex-specific manner.

To investigate the consequences, if any, of skin incision injury on gene expression, we used RNAseq to probe the DRGs and meninges in males and females. As the region of the skin injury is innervated by nerves with cell bodies in DRGs L2-L3 whereas the region of evoked hypersensitivity is innervated by DRGs L4-L5, we investigated transcriptomes in DRGs L2-L3 and DRGs L4-L5. There were 8 DEGs in the DRG L2 and L3 of rats with skin incision compared to naïve rats. However, there were no DEGs in DRG L4 and L5 of rats with skin incision compared to naïve rats. All of the DEGs in DRG L2-3 were increased in expression after skin injury. Of these, *Atf3*, *Gal*, *Sprr1a* and *Csrp3* are all expressed in DRG neurons and are known to increase in response to axonal damage or injury ^39–42^. For example, *Atf3* is the gene that codes for activating transcription factor 3. ATF3 is rapidly induced in response to trauma or cellular stress and plays a key role in axonal regeneration ^40^. *Gal* gene, which codes for Galanin protein, increases in expression in sensory neurons after injury ^39^. *Sprr1a*, which codes for small protein-rich repeat protein 1A, is expressed in DRG neurons after injury, but cannot be detected in naïve DRGs ^42^. Furthermore, after axonal injury *Csrp3* expression increases in the nociceptors in DRG ^41^. As the skin injury inevitably resulted in cutting of a small number of terminal branches of some primary afferents, and as the fold changes of the DEGs we observed was much smaller than that caused by injury to large nerve, it is likely that the transcriptional changes in DRGs L2-3 were primarily the result of damage to a few afferents.

In contrast to the few transcriptional changes in the DRGs, we found surprisingly that skin injury caused substantial transcriptional changes in the meninges in males, but had no effect in females. These observations are in line with our findings from behaviour experiments, where depleting meningeal macrophages unmasked mechanical hypersensitivity in male, but not female, rats with skin incision injury. In males we found that expression of many genes was increased while that of others was decreased. Notably, a number of genes implicated, in studies by others, in driving pain hypersensitivity following inflammation or peripheral nerve injury were decreased in male meninges after skin incision. One of these was *Mmp9* which has been implicated in promoting pain hypersensitivity: after spinal nerve ligation, there is a rapid, but transient upregulation of *Mmp9* gene expression in the affected DRG ^43^. Furthermore, MMP9 (matrix metalloprotease 9) inhibition or deletion attenuates neuropathic pain in the early phase of its development after nerve injury. An additional downregulated gene was *Spp1*, which is implicated in modulating pain processing as inhibiting this gene in male mice with inflammatory pain results in a reversal of sensitivity to both mechanical and thermal stimuli ^44^. Also, *Cxcr4* also contributes to the development and maintenance of hypersensitivity after nerve injury ^45^, and its expression was decreased in male meninges after skin incision injury in this study. Although, *Il1r*, which codes for the protein IL-1 receptor, mediates hypersensitivity in inflammatory models of pain ^46^, is higher in male rats with skin incision injury. From these findings together, we suggest that the transcriptional changes in the meninges after skin injury may be a substrate contributing to the masking of hypersensitivity in male rats.

Overall, we have identified an unsuspected phenomenon whereby, in males, a superficial injury initiates transcriptome reprogramming in the lumbar meninges that we suspect drives an active process, dependent upon CD206^+^ macrophages, which limits the extent of injury-induced pain hypersensitivity. By contrast, in females skin injury produces neither a transcriptome response in the lumbar meninges nor a CD206^+^ macrophage-dependent protection of spread of hypersensitivity. Our findings open up questions as to the mediators and mechanisms by which the effects are mediated in males, and the basis of the profound sex differences. Importantly for the large number of studies of neuropathic pain hypersensitivity produced by sciatic nerve trauma, the skin incision we used was comparable to, or even less extensive than, that used in ‘sham’ controls. Our results demonstrate that, at least in males, this ‘control’ evokes a dramatic response of cells in the lumbar meninges at least one population of which, the CD206^+^ macrophages, we show regulates pain hypersensitivity. These findings have, therefore, potentially broad implications for re-interpreting results from much previous neuropathic pain research.

## Acknowledgments

The authors thank Janice Hicks for assistance throughout this work. We thank Dr. Graham Pitcher for assistance with some of the intrathecal experiments.

## Author contributions statement

MK and MWS conceptualized and designed the experiments. MK performed the experiments; SG contributed to the gene expression analyses; MM contributed to the intrathecal injections; EG contributed to writing and editing; MWS supervised the research; MK and MWS wrote the manuscript.

## Data Availability statement

Data available on request from the authors

