## Supplementary Tables for "Meningeal macrophages mask incision pain sensitization in male rats"

Supplementary Materials

Supplementary Table 1. Differentially expressed genes in the lumbar meninges of male rats with skin incision injury relative to naïve male rats.

| Upregulated genes | logFC | adj.P.Val | Downregulated genes | logFC | adj.P.Val |
|---|---|---|---|---|---|
| Foxf2 | 2.96 | 0.0452 | Mepe | -4.33 | 0.0121 |
| Nos3 | 2.50 | 0.00193 | LOC103690326 | -2.95 | 0.0204 |
| Cldn5 | 2.43 | 0.0111 | Rnase1l1 | -2.86 | 0.00193 |
| LOC120102333 | 2.26 | 0.0204 | Rnase1 | -2.62 | 0.0111 |
| Sox17 (Y) | 2.13 | 0.00528 | Dmp1 | -2.20 | 0.0111 |
| Lypd6 | 1.91 | 0.0407 | Mmp9 | -2.02 | 0.0243 |
| Pappa2 | 1.45 | 0.0423 | Wif1 | -1.79 | 0.00528 |
| Mmrn2 | 1.43 | 0.0129 | Spp1 | -1.60 | 0.00528 |
| Pde3a | 1.39 | 0.00329 | Ifitm6 | -1.38 | 0.0357 |
| Ptprb | 1.28 | 0.0357 | Cxcr4 | -1.01 | 0.0357 |
| Rasip1 | 1.28 | 0.0343 | | | |
| LOC100909737 | 1.28 | 0.0407 | | | |
| Il1r1 | 1.14 | 0.0357 | | | |
| Lama5 | 1.11 | 0.03567 | | | |

Supplementary Table 2. Differentially expressed genes in the DRG L2-L3 of rats with skin incision injury relative to naïve rats.

| Upregulated Genes | logFC | adj.P.Val |
|---|---|---|
| Crisp3 | 5.0 | 0.009 |
| Csrp3 | 4.6 | 0.006 |
| Sprr1a | 4.2 | 0.006 |
| Ucn | 3.3 | 0.006 |
| Mroh4 | 3.0 | 0.009 |
| LOC102551701 | 3.0 | 0.009 |
| Gal | 1.8 | 0.014 |
| Atf3 | 1.41 | 0.04 |
